# Exploring the diversity and physiological characteristics of RubisCO-mediated carbon fixation in culturable prokaryotes

**DOI:** 10.1101/2025.05.01.651632

**Authors:** Arisa Nishihara, Shingo Kato, Moriya Ohkuma

**Affiliations:** Japan Collection of Microorganisms (JCM), RIKEN BioResource Research Center, 3-1-1 Koyadai, Tsukuba, Ibaraki 305-0074, Japan

**Keywords:** CBB cycle, RubisCO, autotroph, metabolic prediction, photosynthesis

## Abstract

The utilization of microbial resources requires their relevant reproducible characteristics, and genome analysis plays a crucial role in discovering valuable strains for future applications. In this study, we analyzed potential carbon-fixing microorganisms via Calvin-Benson-Bassham (CBB) cycle using 6,262 bacterial and 487 archaeal genomes from available cultures in Japan Collection of Microorganisms (JCM), one of the well-established culture collections today. A total of 306 strains (147 genera, eight phyla) carried CBB cycle genes and a literal survey showed that 74 genera had reported evidence of their autotrophic growth, although 73 lacked supporting information. Phylogenetic analysis of RubisCO large subunit (RbcL) identified diverse forms (IA, IB, IC, IE, I+α, II, and III) with distinct metabolic associations: form IA associated with sulfur oxidation and form IC with hydrogen oxidation. Genome-based metabolic predictions suggested potential carbon fixation in numerous strains lacking experimental evidence. Our analyses showed members of *Actinomycetota* harboring form IE RubisCO tend to associate with hydrogen oxidation possibly using oxygen or nitrate as an electron acceptor. Additionally, 12 strains in *Pseudomonadota* contained *pufL* and *pufM* genes, suggesting possible phototrophic capabilities, although some failed to predict their electron donors and they possibly use CBB cycle to regulate intracellular redox balance under photoheterotrophic growth. Our findings highlight unrecognized autotrophic potentials in JCM strains and expand our knowledge of carbon fixation diversity. Future experimental validation will deepen our understanding of these microbes’ roles in the global carbon cycle, with potential applications in carbon sequestration and environmental sustainability.

## Introduction

The Calvin-Benson-Bassham (CBB) cycle is a fundamental biochemical pathway for carbon fixation, playing a pivotal role in the global carbon cycle. Today, other six natural carbon fixation pathways converting atmospheric CO_2_ to organic carbon have been identified in the autotrophic microorganisms; the reverse tricarboxylic acid (rTCA) cycle including a variant [reversed oxidative TCA cycle (roTCA)], the 3-hydroxypropionate (3HP) bi-cycle, the 4-hydroxybutyrate/3-hydroxypropionate (4HB/3HP) cycle, the dicarboxylate/4-hydroxybutyrate (DC/4HB) cycle, the reductive acetyl-CoA (Wood–Ljungdahl) pathway, and the reductive glycine pathway. Among these pathways, the CBB cycle is found to distribute in photosynthetic organisms not only bacteria but also Eukaryotes, such as higher plants and eukaryotic alga, and in various chemolithoautotrophic bacteria. It is the primary carbon fixation mechanism, accounting for ∼95% of the carbon fixed on Earth (Field *et al*., 1998), which is catalyzed by a pivotal enzyme, Ribulose-1,5-bisphosphate carboxylase/oxygenase (RubisCO) (Banda *et al*., 2020; Schulz *et al*., 2022).

The RubisCO family has been classified into multiple structural and functional forms, including forms I, II, III, and IV, which differ in their catalytic efficiency, substrate specificity, and evolutionary history (Prywes *et al*., 2023; Tabita *et al*., 2008). With the exception of form IV RubisCOs (RubisCO-like proteins), which catalyze ribulose-1,5-bisphosphate carboxylation, all RubisCO forms have been reported to be involved in CO_2_ fixation. Form II RubisCO is characterized by a poor affinity for CO_2_, indicating the adapted function in low-O_2_ and high-CO_2_ environments (Badger and Bek, 2008). Since the RubisCO has oxygenase activity that competes with carbon dioxide fixation, form I RubisCOs have recruited the small subunit, whose encoding gene (*rbcS*) usually colocalizes in an operon with the large subunit gene (*rbcL*), which increases the specificity for CO_2_ in the ancient era before oxygen was abundant on the Earth (Nishihara *et al*., 2024; Schulz *et al*., 2022). The form I RubisCOs have been further classified into subgroups: IA and IC (mostly found in *Pseudomonadota*), IB (*Cyanobacteria*, green alga and plants), ID (eukaryotic alga), IE (*Actinomycetota*), and *Thermus* group (Schulz *et al*., 2022). Recently, based on *rbcL* phylogeny, four deeper branching orthologs of form-I RubisCO, which do not have *rbcS*, have been identified from metagenome-assembled genomes, expecting a key to elucidate the small subunit innovation for increasing the CO_2_ affinity, and they are designated as forms I’(Banda *et al*., 2020), I-Anaero (Schulz *et al*., 2022), I-α and I’’ (West-Roberts *et al*., 2021), respectively. Form III RubisCO lacking phosphoribulokinase (PRK) has been reported to be associated with the pentose bisphosphate pathway found in some methanogenic archaea (Aono *et al*., 2015; Aono *et al*., 2012; Sato *et al*., 2007). However, autotrophy via this pathway remains uncertain, as methanogens primarily rely on the Wood–Ljungdahl pathway for carbon fixation (Kono *et al*., 2017). More recently, form III RubisCO harboring PRK was reported in a chemolithoautotrophic bacterium, *Thermodesulfobium acidiphilum* (*Thermodesulfobiota*), suggesting the possible involvement of a modified CBB cycle for CO_2_ fixation (Frolov *et al*., 2019). These reports provide variations in the function of CBB associated genes in microbial metabolism.

In phototrophic bacteria, all purple sulfur bacteria have been reported to fix CO_2_ via the CBB cycle, which belongs to *Gammaproteobacteria*. Purple non-sulfur bacteria have been found in *Alphaproteobacteria* and *Betaproteobacteria*, some of which could fix CO_2_ via the CBB cycle but grow well under heterotrophic conditions (Imhoff, 2009). In purple non-sulfur bacteria, RubisCO-mediated carbon assimilation is known to contribute to the alleviation of an over-reduced state (Shimada and Takaichi, 2024). Some strains have been reported to possess both forms I and II, different clusters of form I, or only form I or II. Aerobic anoxygenic phototrophic bacteria (AAPB), which are widely distributed in *Alpha-*, *Beta*-, and *Gammaproteobacteria*, are obligately aerobic and show heterotrophic growth, and they have not yet demonstrated autotrophic growth (Yurkov and Beatty, 1998). However, light-stimulated CO_2_ uptake has been reported in them, such as *Acidiphilium rubrum* (*Acetobacterales*, *Alphaproteobacteria*) (Kishimoto *et al*., 1995) and *Erythrobacter longus* (*Sphingomonadales, Alphaproteobacteria*) (Shiba, 1984; Shiba and Harashima, 1986; Yurkov and Beatty, 1998). In AAPB, the CO₂ fixation through the CBB cycle is insufficient to sustain autotrophic growth but may contribute to the supply of additional organic carbon intermediates for heterotrophic growth (Yurkov and Beatty, 1998). Therefore, autotrophic growth via the CBB cycle does not always correspond to the presence of functional genes. A comprehensive investigation into the involvement of CBB in the various strains requires a thorough examination of their cultivation, with particular attention to the uptake of CO₂ and/or CO₂-independent growth.

To our knowledge, few studies are focusing on the connection of both genomic information and physiological characteristics through the culture-based data targeting diverse phylogeny with large-scale genome analysis, despite the increase of identification of autotrophic bacteria through genome and/or metagenome studies (Garritano *et al*., 2022). The Japan Collection of Microorganisms (JCM), one of the well-established culture collections in the world today, maintains cultivable strains for reproducible experiments with well-characterized type strains as well as non-type strains as other culture collections (*e.g.*, Leibniz Institute DSMZ, German Collection of Microorganisms and Cell Cultures). In this study, we analyzed potential carbon-fixing JCM strains that harbored the CBB cycle genes, identifying the forms of RubisCO and connecting with information from reported culture-based data as possible, and predicted their metabolic potentials based on genome information for the strains that have not demonstrated autotrophic growth yet. Our findings provide a foundation for future research aimed to harness microbial resources for basic studies and their biotechnological applications, including carbon sequestration and sustainable environmental management.

## Materials and Methods

### Annotation of the genomes for the CBB cycle

All genomes used in this study were collected from the RefSeq database in National Center for Biotechnology Information (NCBI) (https://www.ncbi.nlm.nih.gov). To evaluate the capacity for the CBB cycle, the collected genomes were annotated using METABOLIC (v4.0) against the implemented HMM databases (KEGG KOfam, Pfam, TIGRfam, and custom HMMs) with the default options (Zhou *et al*., 2022). According to the results of the METABOLIC annotation results, we selected organisms as a possibly CBB cycle harboring bacteria that possessed all four selected KEGG module step hits – M00165+01 [K00855 (PRK, *prkB*)], M00165+02 [K01601 (*rbcL*, *cbbL*)], M00165+03 [K00927 (PGK)], and M00165+04 [K05298 (GAPA), K00150 (*gap2*, *gapB*), or K00134 (GAPDH, *gapA*)].

### Prediction of the metabolic potential of putative autotrophs

Using strains predicted to be involved in the CBB cycle, the cultivation conditions for carbon fixation were examined for each representative species within the genera through a literature survey (as of December 2024) of all available publications on Google Scholar (https://scholar.google.co.jp). If no report of autotrophic growth was found within a genus, the metabolic potential for autotrophic growth was predicted based on the presence of functional proteins, as annotated by METABOLIC, with reference to the MetaCyc database (Karp *et al*., 2019) (Table S1a). The prediction were focused on proteins associated with electron acceptors (oxygen availability or anaerobic respiration using sulfur species and nitrate) and electron donors (hydrogen and sulfur species), as listed in Table S1b. Additionally, photosynthesis-related genes for type-II reaction center proteins (*pufM* and *pufL*) were identified through a BLAST homology search using biochemically characterized proteins registered in SwissProtKB (Boutet *et al*., 2007), after all predicted protein-coding genes of the representative species were annotated using Prokka (v1.14) (Seemann, 2014) (Table S1c).

### Phylogenetic analysis of rbcL gene

The gene of *rbcL* were collected from all predicted protein-coding gene sets from possibly autotrophic JCM strains. For the construction of a phylogenetic tree, representative sequences were selected by clustering with CD-HIT v.4.8.1 (Fu *et al*., 2012) at an 80% identity threshold. Clustering was performed among representative sequences of the same genus with no documented evidence of autotrophic growth based on a literature survey. A sequence alignment for the selected sequences was generated using MAFFT (v7.427) with the options, --localpair and --maxiterate 1000 (Katoh *et al*., 2005). The alignment was subsequently trimmed using trimAl (v1.4.rev22) (Capella-Gutiérrez *et al*., 2009) with a gap threshold of 0.8. The final alignment was then used as input for maximum-likelihood phylogenetic estimation via IQ-TREE (v2.1.3) (Nguyen *et al*., 2014). The tree was constructed using the profile mixture model, which was identified as the best-fit model by ModelFinder (Kalyaanamoorthy *et al*., 2017), implemented in IQ-TREE (v2.1.3) and 1,000 ultrafast bootstrap replicates. Transfer bootstrap support values were calculated using BOOSTER (v0.1.2) with 1000 replicates from ultrafast bootstrap (1000 replicates) (Lartillot and Philippe, 2006).

## Results and Discussion

### Carbon fixation ability of strains harboring CBB cycle gene

In this study, we analyzed the genomes of 6,262 bacterial and 487 archaeal strains distributed across 1,776 genera within 40 phyla, which is available in the JCM culture collection. Among these genomes, 306 strains (147 genera within eight phyla) were found to be potentially involved in the CBB cycle based on the presence of selected prodigal modules annotated using the METABOLIC tool (Zhou *et al*., 2022), as listed in Table S2. Most of them were found in *Pseudomonadota* (207 strains), followed by *Actinomycetota* (82 strains), and as for archaea, only six strains belonging to *Methanobacteriota* were identified.

We then conducted a literature survey to determine whether carbon fixation abilities have been reported within these genera. The strains used for literature-based surveys are listed in Table S2 as representative strains. Among the 147 genera, 74 contained any species with reported carbon fixation capabilities but 73 lacked supporting information. The CBB-cycle gene harboring JCM strains that reportedly showed carbon fixation belonged to 54 genera. In addition, the carbon fixation conditions of these strains, such as electron donors and acceptors, were examined as described in Tables S3 and summarized in Table 1 (discuss later). These genera that have been reported autotrophic growth were distributed across five phyla: *Actinomycetota*, *Thermodesulfobiota*, *Pseudomonadota*, *Thermodesulfobacteriota*, and *Methanobacteriota* (Table 1, Table S3). Conversely, as far as we investigated, 73 genera were not associated with any reports of autotrophic growth; 174 species had literally no reports of autotrophic growth, including 15 species that have been reported as lacking autotrophic growth.

**Table 1.**
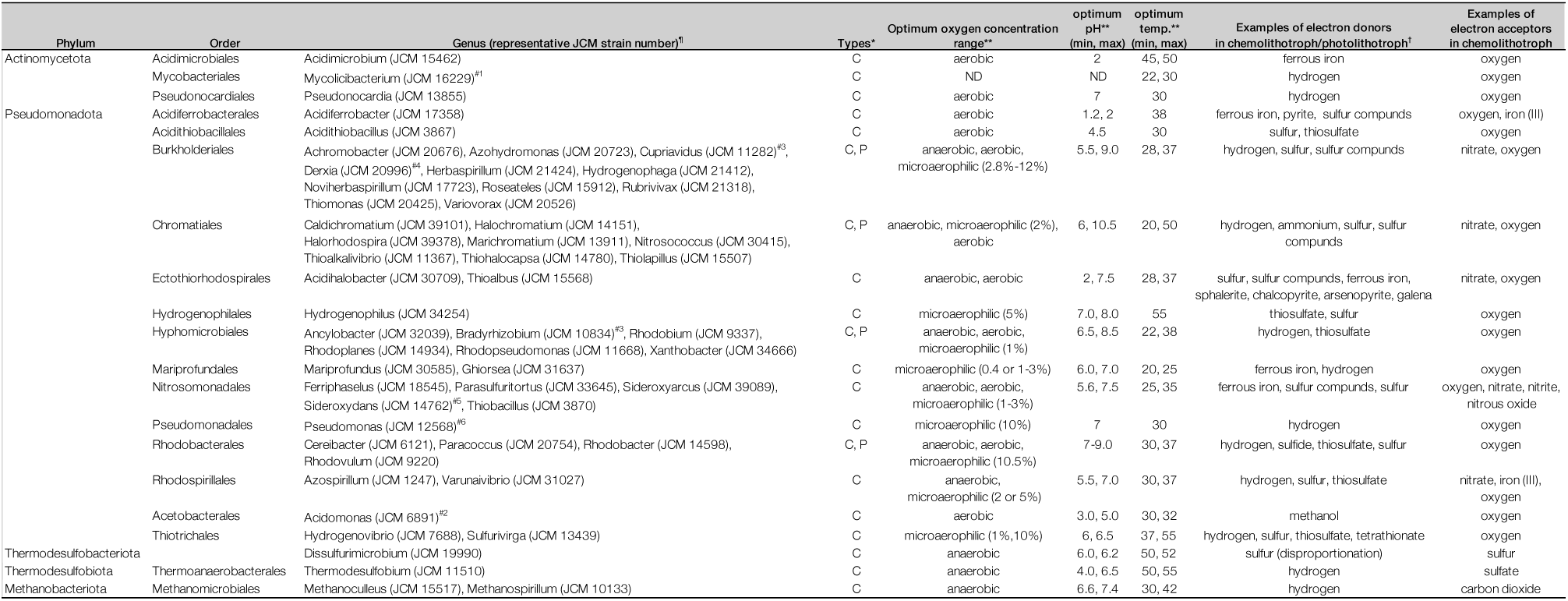
Summary of autotrophs and representative growth conditions for each genus in the JCM culture collection. ^¶^, Representative strains are indicated in Table S2. Full strain names and corresponding data are provided in Table S3. *, Autotrophic growth under chemolithotrophic (C) or phototrophic (P) conditions. **Data on optimum or tested oxygen concentration under autotrophic conditions, temperature, or pH were extracted from the reference; ND indicates “Not Determined” (could not be found in the literature survey). ^†^, Electron donors comprising various reduced sulfur compounds (e.g., sulfide, thiosulfate, tetrathionate) are collectively referred to as sulfur compounds. ^#1^ Autotrophic growth was not observed under conditions with CO and O₂ in the gas phase. ^#2^ Initially, growth depended on autotrophic carbon fixation, and carbon assimilation was subsequently coupled with the ribulose monophosphate (RuMP) pathway during the early to mid-log phases of methylotrophic growth. ^#3^ The strain used in the experiment demonstrating autotrophic growth is different from the JCM strain. ^#4^ Source information not available. ^#5^ Unvalidated species. ^#6^ Autotrophic growth via CO oxidation has been reported instead of CO_2_ fixation.

Based on the literature survey, within *Actinomycetota*, only three genera [*Acidimicrobium* (*Acidimicrobiales*), *Mycolicibacterium* (*Mycobacteriales*), *Pseudonocardia* (*Pseudonocardiales*)] contained strains that have been reported to exhibit autotrophic growth, despite 21 genera being annotated as putative autotrophic bacteria based on genome information (Table 1, S2, and S3). All the species demonstrating autotrophic growth in these three genera were aerobic chemolithoautotrophs, utilizing hydrogen or ferrous iron as electron donors and oxygen as the electron acceptor (Table 1 and S3).

*Pseudomonadota* exhibited the most diverse range of metabolic types, including photosynthetic organisms (*e.g*., purple sulfur or non-sulfur bacteria), reflecting the wide phylogenetic diversity within this phylum. Photoautotrophs were identified in the classes *Alphaproteobacteria* (*Rhodoplanes*, *Rhodopseudomonas*, and *Rhodobium* in *Hyphomicrobiales* and *Cereibacter*, *Rhodobacter*, and *Rhodovulum* in *Rhodobacterales*), *Betaproteobacteria* (*Rubrivivax* in *Burkholderiales*), and *Gammaproteobacteria* (*Caldichromatium*, *Halochromatium*, *Marichromatium*, *Thiohalocapsa*, and *Halorhodospira* in *Chromatiales*), and they reportedly utilized hydrogen or sulfur compounds (*e.g.*, thiosulfate and sulfide) as electron donors (Table 1 and S3). Among chemolithoautotrophs, microaerophilic conditions (0.4–5% oxygen) were required for members of the classes “Zetaproteobacteria” (*Mariprofundus* and *Ghiorsea* in *Mariprofundales*) and *Hydrogenophilia* (*Hydrogenophilus* in *Hydrogenophilales*). Members of the class *Alphaproteobacteria* (*Acetobacterales, Hyphomicrobiales*, *Rhodobacterales*, and *Rhodospirillales*) required microaerophilic (1–10.5% oxygen) or aerobic conditions for chemolithoautotrophy. Most bacteria within the classes *Betaproteobacteria* (*Burkholderiales* and *Nitrosomonadales*) and *Gammaproteobacteria* (*Acidiferrobacterales*, *Chromatiales*, *Ectothiorhodospirales*, *Pseudomonadales*, and *Thiotrichales*) also required oxygen (ranging from microaerophilic to aerobic) but most species in *Betaproteobacteria* specifically required microaerophilic conditions. Furthermore, some species demonstrated autotrophic growth only under anaerobic conditions, utilizing nitrate as an electron acceptor such as three species in *Noviherbaspirillum*, *Rubrivivax*, and *Thiomonas* in *Burkholderiales* (Table 1 and S3).

Species in other phyla, *Thermodesulfobiota* (*Thermodesulfobium narugense*), *Methanobacteriota* (*Methanoculleus horonobensis* and *Methanospirillum hungatei*), and *Thermodesulfobacteriota* (*Dissulfurimicrobium hydrothermale*), were reported to have autotrophic capacity under anaerobic conditions (Ferry *et al*., 1974; Mori *et al*., 2003; Shimizu *et al*., 2013; Slobodkin *et al*., 2016). Since hydrogenotrophic methanogenic *Methanobacteriota* are generally known to fix carbon through methanogenesis via the Wood-Ljungdahl (WL) pathway (Borrel *et al*., 2016), CBB cycle is not mainly used for their autotrophy. Instead, the sulfur metabolism-related bacteria listed in Table 1 seem to be involved in CO_2_ fixation via their CBB cycle. In the reported culture conditions demonstrating their autotrophic growth, *T. narugense* was associated with sulfate-reducing conditions coupled with hydrogen oxidation, while *D. hydrothermale* was associated with the metabolism of elemental sulfur disproportionation. Since autotrophic growth of sulfate-reducing bacteria has been reported in one of the members in the *Thermodesulfobium* (*T. acidiphilum*), which was found to fix CO_2_ via the CBB cycle (Frolov *et al*., 2019), *T. narugense* may also possess the capability to fix CO₂ through a similar mechanism. In *D. hydrothermale*, a complete gene set for WL pathway has previously been reported in the genome (Yvenou *et al*., 2021). While a previous study suggested that the CBB cycle gene set was incomplete, our analysis indicates that *D. hydrothermale* possesses a complete set of CBB cycle genes based on annotations from KEGG (GhostKOALA) (Kanehisa *et al*., 2016) (see later discussion regarding the form I-α RbcL).

### RubisCO enzyme forms and physiological characteristics for autotrophy

To explore the types of RubisCO found in JCM strains and their associated metabolisms, RubisCO large subunit sequences (RbcL) were classified based on phylogenetic clustering analysis, and metabolisms related to autotrophic growth are shown in Fig.1. In the analysis, form IV RubisCOs (RubisCO-like proteins) and archaeal PRK-lacking form III RubisCOs which are associated with the pentose bisphosphate pathway were omitted (Aono *et al*., 2015; Aono *et al*., 2012; Sato *et al*., 2007; Tabita *et al*., 2008). Various RubisCO forms (IA, IB, IC, IE, I+α, II, and III) were identified across multiple JCM strains. In our constructed phylogeny, both the reduced phylogeny (Fig. 1 and Fig S1) and the phylogeny including all sequences (Fig. S2) showed the same topology – the form I-Anaero clade was located on a deeper branch of the form IA/IB groups, positioned as a sister group to the form IC/IE and *Thermus* clade. In a recent study, it was mentioned that the form I-Anaero clade is unstable in phylogenetic reconstructions; its placement is different from previous studies, but our constructed tree has the same topology as that of the reported RbsS (Liu *et al*., 2023; Schulz *et al*., 2022). Therefore, we focused on annotating its phylogenetic distribution based on Fig.1 rather than discussing the tree topology.

**Figure. 1.**
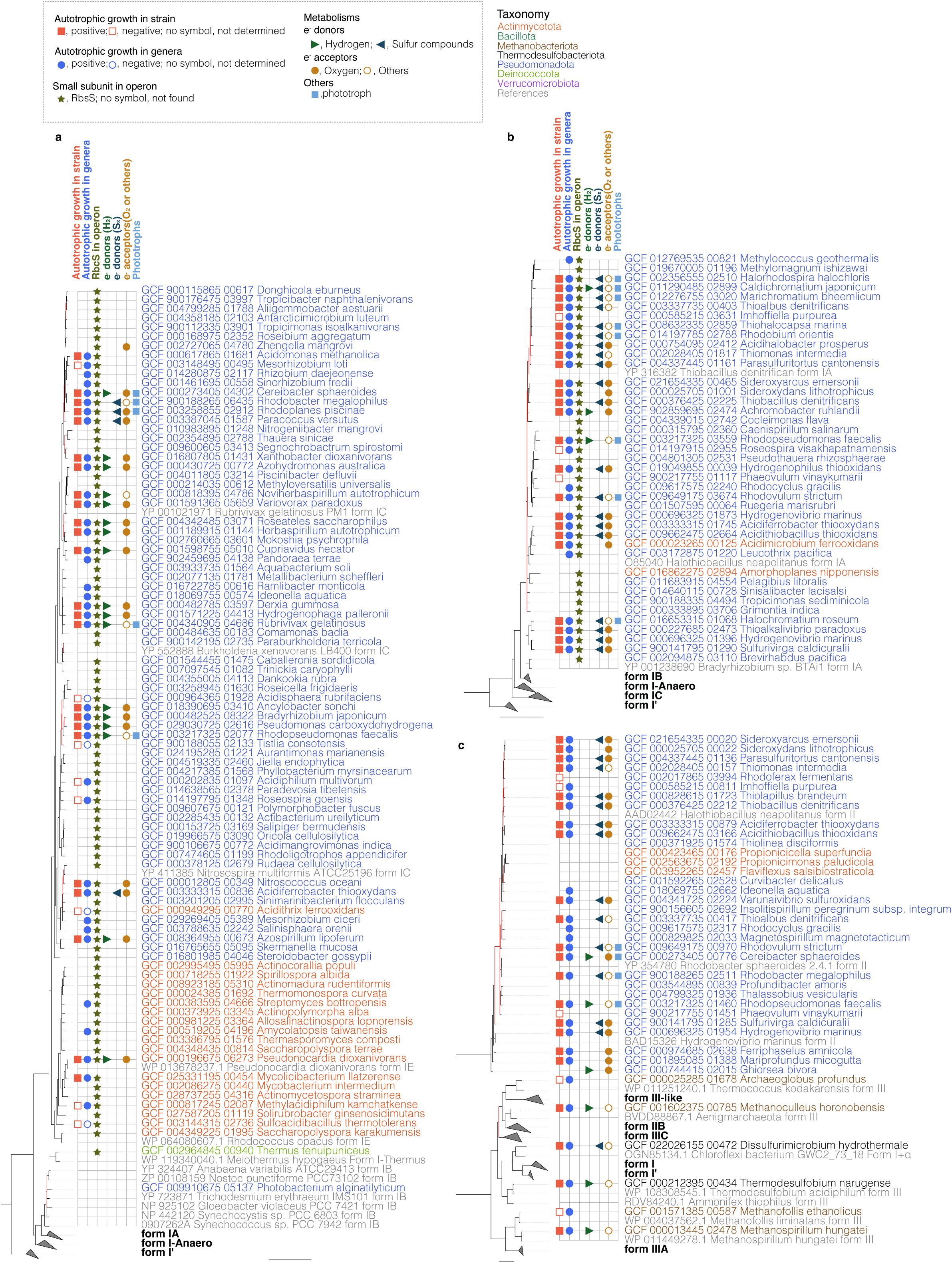
A maximum likelihood tree of the amino acid sequences of RbcL. Species with confirmed carbon fixation were included in the phylogenetic tree as representatives of their respective genera. Constructed phylogenetic trees are shown without grouping branches of (a) form IC, (b) form IA, and (c) form II RbcL. Reference sequences were collected from published data (Badger and Bek, 2008; Prywes *et al*., 2023; Schulz *et al*., 2022). The tree was constructed using IQ-TREE with the LG+C40+G4+F substitution model and 1,000 ultrafast bootstrap replicates. Input sequences were trimmed using trimAl (-gt 0.8), resulting in 211 sequences with 438 amino acid positions. Bootstrap branch support values were calculated using BOOSTER (v0.1.2) from ultrafast bootstrap (1000 replicates). Unsupported branches with lower bootstrap values (≤90%) were collapsed, and branches with bootstrap values <95% were colored red. The scale bar indicates 1.0 changes per amino acid site. For each strain, the presence of reported metabolic activity associated with autotrophic growth is shown, based on data provided in Table S3. See Fig. S1 for the full tree.

In total, form IA RbcL was identified in 42 strains, of which 23 exhibited carbon fixation capabilities. Form IB was previously found only in a *Cyanobacteria* in Prokaryotes but we found one strain, *Photobacterium alginatilyticum* (*Pseudomonadota*), in this clade, which indicated that the strain obtained *rbcL* via horizontal gene transfer from *Cyanobacteria*. Form IC contained the most abundant 97 strains, 22 of which demonstrated carbon fixation capabilities. Form IE, associated with the *Actinomycetota* cluster, was identified in 61 strains; however, only 2 strains exhibited carbon fixation abilities. Although RbcL and other genes associated with the CBB cycle have been reported to be absent in some species (Müller *et al*., 2016), all eight JCM strains of *Thermus* possessed the RbcL gene belonging to form I *Thermus* group, consistent with previous reports for some members of the genus (Müller *et al*., 2016; Schulz *et al*., 2022). To our knowledge, only heterotrophic growth has been reported with no autotrophic demonstration in the genus, although there are reports of the existence of RubisCO in the genome. Form II was identified in 33 strains, including 19 carbon fixation reported strains. As for form III RbcL, only one species, *T. narugense* (*Thermodesulfobiota*), was found, and its RbcL gene showed 94.87% amino acid sequence similarity to that of *T. acidiphilum*, which has been demonstrated to fix CO_2_ through a RubisCO-mediated transaldolase variant of the CBB cycle. Since *T. narugense* possesses a complete gene set for the CBB cycle, as described in the previous section, it may also be capable of catalyzing CO₂ fixation as *T. acidiphilum* (Frolov *et al*., 2019).

Strains that contained RbcL of forms IA, IC, and II, mainly composed of the members of *Pseudomonadota.* Six strains from *Actinomycetota* were grouped with *Pseudomonadota* in the later-diverging phylogenetic branch, suggesting that some of these members may have acquired *rbcL* via horizontal gene transfer from *Pseudomonadota*. From those *Actinomycetota*, *Acidithrix ferrooxidans* (form IC RbcL), the only species described in the genus, has been reported that it does not have the ability to fix CO_2_ (Jones and Johnson, 2015). In contrast, *Acidimicrobium ferrooxidans* (form IA RbcL) has reported the autotrophic growth (Norris, 1996). Three orthologs of form I RbcL without RbcS have been identified from metagenome-assembled genomes: I’(Banda *et al*., 2020), I-Anaero (Schulz *et al*., 2022), I-α (West-Roberts *et al*., 2021), respectively. One JCM strain of *D. hydrothermale* was found to belong to the form I-α clade, which does not possess the *rbcS* gene based on a BLAST search of its complete genome (GCF_022026155). The finding of RbcL in the culturable autotrophic strain of *D. hydrothermale* is intriguing, and it is possibly an ancient form of form I RubisCO working without the small subunit. Further research on elucidating the mechanisms of these enzymes is expected to contribute to understanding of the evolution of form I RbcL with the development of the small subunit. Besides, *P. alginatilyticum* (*Pseudomonadota*) with RbcL classified as form IB doesn’t possess RcbS but it apparently derived from horizontal gene transfer from *Cyanobacteria* (see above).

As for the association of metabolic diversity and RubisCO forms, we examined metabolic information of autotrophic growth based on literature surveys focusing on electron donors (hydrogen or sulfur species) and acceptors and the ability of photoautotrophs. Utilization of hydrogen as electron donors is abundant in strains harboring form IC RbcL, accounting for 10 of 19 strains (Fig.1 and Table S3). In contrast, strains utilizing sulfur compounds (*e.g.*, sulfur, thiosulfate, and S°) were more abundant in form IA, accounting for 10 of 11 strains. Strains showing phototrophic ability were mostly found in form IA, accounting for eight of 11 strains. Additionally, although all the strains harboring form II do not have form I (26 strains have form IA and 10 have form IC), all the phototrophs harboring form II also have form IA or IC. These data may suggest that different RubisCO forms could be associated with metabolic variation in electron donor utilization such as hydrogen vs sulfur compounds, although the strains belong to different phylogenetic groups (*e.g*., at the class and order level).

### Metabolic predictions of potential autotrophs

Among the 174 strains that were either found to lack carbon fixation ability or had not yet been examined but which possessed genes involving the CBB cycle in their genomes, 144 belonged to genera that do not report any autotrophic growth demonstrations (Table S2); 76 strains from 54 genera in *Pseudomonadota*, 57 strains from 16 genera in *Actinomycetota* and 7 strains from one genus in *Deinococcota*. Then, we examined their metabolic potentials to predict possible autotrophic conditions, especially focusing on electron acceptors (oxygen availability or anaerobic respiration using sulfur species and nitrate) and donors (hydrogen and sulfur species) (Table S4).

Regarding hydrogen utilization as an electron donor, we explored the potential presence of uptake hydrogenases. In the hydrogenase functional group, hydrogenases annotated as Group A2 and A3 ([FeFe]-hydrogenases) and Group 1, 2a, 2b, 2c, 2d, 2e, 4h, and 4i ([NiFe]-hydrogenases) were selected as uptake hydrogenases, based on the annotation results from METABOLIC (Sondergaard *et al*., 2016; Zhou *et al*., 2022). While [NiFe]-hydrogenases were detected, our analysis did not find the [FeFe]-hydrogenase genes in the strains examined. Nearly half of the members possessed the

[NiFe]-hydrogenase gene, suggesting their potential utilization of hydrogen as an electron donor (Table 2 and Table S5). The prevalence of hydrogen-utilizing bacteria was highly observed in strains harboring form IC RbcL, which is consistent with the results reported for other bacteria as described above. Form IE was grouped with those of *Actinomycetota* and contained fewer bacteria demonstrating autotrophic growth. Among the strains, we found that the number of strains possessing [NiFe]-hydrogenase gene (33 in 53 strains) was higher than that of sulfur-utilization (five in 53 strains). As for form IA, previous studies showed a predominance of sulfur-utilizing bacteria (described above); however, among species in the genera for which carbon fixation has not yet been demonstrated, the number of strains potentially utilizing sulfur vs hydrogen was not much different; nine strains harbored genes for sulfur-utilization and seven had hydrogenase gene in the genome (Table S5). For terminal oxidases, most strains of *Pseudomonadota* that were analyzed possessed key genes of terminal oxidases, both for microaerophilic types, which have high affinity for oxygen [heme‒copper oxidases family (HCOs)-C types (*ccoON*) and cytochrome *bd*-type oxidases (*cydAB*)] and for aerobic types, which have low oxygen affinity [(HCOs-A type terminal oxidase (*coxAB* or *cyoABCD*)] (Morris and Schmidt, 2013). In contrast, other taxa, especially *Bacillota* and *Deinococcota*, tend to possess either of these types (Table 2).

**Table 2.**
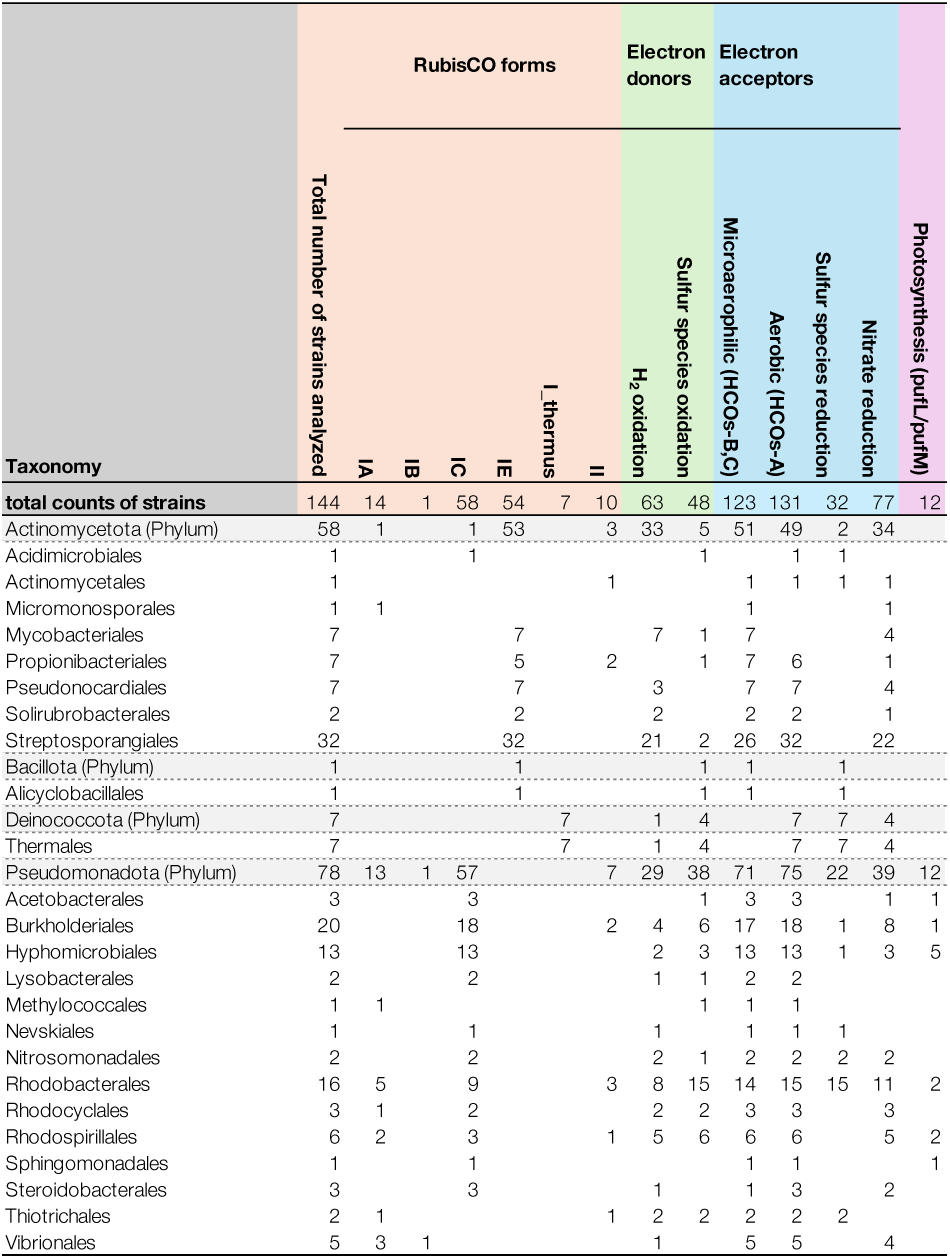
Summary of the strain number corresponding to predicted metabolisms based on genome information. Strains used in this analysis are derived from genera with no reported evidence of autotrophic growth. The light gray highlights indicate the total number for each phylum. Details of the genes used for metabolic predictions are given in Table S1. Full strain names and corresponding data are provided in Table S4.

In *Pseudomonadota*, the highest number of strains was found in *Paraburkholderia* (*Burkholderiales*) with nine strains, followed by *Jiella* (*Hyphomicrobiales*) with four strains. The remaining genera were found to contain three or fewer strains. In the phylum, it seems that hydrogen and sulfur species were mainly used as electron donors and oxygen seems to be used as an electron acceptor under chemolithoautotrophic conditions. A diverse range of strains have the potential to fix carbon through CBB cycle, including 11 species that belong to each mono-typic genus in the classes *Alphaproteobacteria* [*i.e*., *Tistlia* (form IA), *Zhengella* (form IC), *Segnochrobactrum* (form IC), *Dankookia* (form IC), *Profundibacter* (form II), *Thalassobius* (form II), *Insolitispirillum* (form II)] and *Gammaproteobacteria* [*i.e*., *Methylomagnum* (form IA), *Cocleimonas* (form IA), *Met allibacterium* (form IC), *Rudaea* (form IC)] (Table S5).

In *Actinomycetota*, strains in *Flaviflexus* (*Actinomycet ales*), *Amorphoplanes* (*Micromonosporales*), *Propionicicella*, and *Propionicimonas* (*Propionibacteriales*) did not possess form IE RbcL, but did either form IA or II, while all the others in this phylum had form IE (Table S5). The genera containing the most observed potential autotrophic strains within this phylum were *Actinomadura* (*Streptosporangiales*) with 24 strains, followed by *Mycobacterium* (*Mycobacteriales*) with seven strains, and *Actinomycetospora* (*Pseudonocardiales*) and *Thermomonospora* (*Streptosporangiales*) with five strains, respectively. In *Actinomadura*, all strains in this genus possessed the gene for low-affinity terminal oxidase [HCOs-A (*coxA*/*coxB*)] and 18 strains additionally had the genes for high-affinity terminal oxidase, cytochrome *bd*-type oxidases (*cydA*/*cydB*), suggesting the ability to grow under microaerophilic to aerobic conditions. The thiosulfate oxidation gene (*sqr*) was found in two strains that possessed cytochrome *bd*-type oxidases (*cydA*/*cydB*), and they possibly grow autotrophically with thiosulfate as electron donor under microaerophilic to aerophilic conditions. [NiFe]-hydrogenase gene was annotated in 15 strains with the existence of nitrate reduction genes (*narG*/*narH*) in 12 strains, supposing the possibility of CO_2_ fixation with hydrogen-oxidization using hydrogen as an electron donor and oxygen as an electron acceptor under aerobic conditions or with nitrate reduction using hydrogen as an electron donor and nitrate as an electron acceptor. The same metabolisms are thought to be involved in strains of *Actinomycetospora* and *Thermomonospora*, since all the strains in these two genera possessed the gene for high-affinity terminal oxidase and almost half of them possessed the genes for [NiFe]-hydrogenase and nitrate reduction (*narG*/*narH*).

As for *Thermus*, it consists of thermophilic strains that have been isolated worldwide in geothermal environments (≥55°C) ranging from pH 5.0 to 10.5. These strains have been demonstrated heterotrophic growth, although some have been reported to harbor complete gene sets for CBB cycle (Da Costa *et al*., 2006; Müller *et al*., 2016). All RbcL sequences from *Thermus* grouped in the same cluster as shown in the previous study (Müller *et al*., 2016; Schulz *et al*., 2022), and the genome analysis showed that strains in this genus had the gene for low-affinity terminal oxidase [HCOs-A (*coxA*/*coxB*)]. Some strains of the genus had genes for complete thiosulfate oxidation proteins (*soxABXYZ*), and the other had an incomplete set of genes lacking only *soxB*. In *Thermus sediminis*, chemolithoautotrophic growth was reported under conditions involving the oxidation of thiosulfate to sulfate, although this characteristic has been only described in the abstract, and the details remain uncertain (Zhou *et al*., 2018). Our genome analysis also suggested that if carbon fixation occurs, it is expected to take place under thiosulfate oxidation conditions.

As for the autotrophic potential of phototrophic bacteria, because the presence of CBB cycle genes does not always link to active carbon fixation in phototrophs (Shimada and Takaichi, 2024), we also examined photosynthesis-related genes for further insights into the genera that have not been demonstrated to have autotrophic ability at the genus level. The *pufL* and *pufM* genes, which encode a type II reaction center involved in anoxygenic photosynthesis, were identified in the 12 strains in *Pseudomonadota* through a homologous BLAST search. Those belonged to the seven genera such as *Jiella*, *Aliigemmobacter, Phaeovulum, Dankookia, Skermanella, Polymorphobacter* in *Alphaproteobacteria* and *Piscinibacter* in *Betaproteobacteria*. All of those have the form IC RubisCO except one strain, *Phaeovulum vinaykumarii*, which possessed genes for both Form IA and II RubisCOs; however, it has been reported that it lacked autotrophic growth ability in the photosynthetic conditions (Srinivas *et al*., 2007) although autotrophic growth in *Rhodobacter* and *Rhodovulum* (*Paracoccaceae*, *Rhodobacterales*) has been reported.

In the genus *Jiella* (*Alphaproteobacteria*), four strains were identified as potential phototrophs, representing the highest number of strains found among the genera analyzed. In the genomes of the four strains, genes involved in hydrogen or sulfur compound oxidation were not detected (Table S5), requiring further analysis for predicting the metabolisms of autotrophic growth. Alternatively, it is possible that *Jiella* utilizes the CBB cycle under heterotrophic growth conditions and does not involve autotrophic growth, as reported in aerobic anoxygenic phototrophic bacteria (AAPB)(Yurkov and Beatty, 1998). In *Alphaproteobacteria*, *Hyphomicrobiales* that contain *Jiella* (*Aurantimonadaceae*) and *Rhodoligotrophos* (*Parvibaculaceae*) accommodate multiple photoautotrophic purple non-sulfur bacteria (*e.g.*, *Rhodobium* and *Rhodopseudomonas*) (Table S3) and AAPB (*e.g*., *Methylorubrum*) (Shimada and Takaichi, 2024). In *Acetobacterales*, which includes *Dankookia*, lacking the genes involved in electron donor utilization as *Jiella* (Table S5), has multiple reports for AAPB (*e.g.*, *Acidisphaera* and *Craurococcus*) and purple non-sulfur bacteria (*e.g.*, *Rhodopila* and *Rhodovastum*) (Shimada and Takaichi, 2024).

## Conclusion

This study identified previously unrecognized autotrophic potential via CBB cycles in a diverse accessible JCM strains through an integrated analysis of genome data and literature surveys. We clarified which carbon fixation capabilities have been experimentally demonstrated and what remains unclear regarding the carbon fixation potential of isolated strains. These findings provide a foundation for future research focused on validating autotrophic capability through further experimental analysis.

## Acknowledgments

This work was supported by Japan Science and Technology Agency (JST) GteX program Grant Number JPMJGX23B0 (M.O.) and partially supported by Japan Society for the Promotion of Science Grant-in-Aid for Early-Career Scientists 24K17163 (A.N.).

## Supplemental table legends

**Table S1.** Lists of key genes associated with electron acceptors and electron donors for metabolic prediction. (a) Summary of key gene sets for annotating metabolic potential in this study, (b) annotation details based on Metabolic, and (c) photosynthetic gene used for BLAST search.

**Table S2.** List of microbial strains analyzed in this study.

*, Strains used for literature-based surveys, phylogenetic analysis, and metabolic potential analysis are marked with a “+”.

^†^, For strains whose carbon fixation potential could not be determined from literature data, “ND” (Not Determined) is indicated.

**Table S3.** Autotrophic species and representative growth conditions for each genus in the JCM culture collection. The summary is available in Table 1.

* Data on optimum temperature or pH were extracted from the reference; ND indicates “Not Determined” (could not be found in the literature survey).

**Table S4.** Counts of presence of key gene sets for metabolic prediction and counts of RubisCO forms in the genome. Metabolic predictions were performed based on criteria listed in Table S1a, b, c. Annotation of RubisCO forms is based on Fig.1, Fig. S1, and Fig. S2. A summary of metabolic prediction counts for strains with no reported evidence of autotrophic growth is provided in Table 2.

## Supplemental figure legends

**Figure S1.** A maximum likelihood tree of the amino acid sequences of RbcL. With the exception of the grouping of RbcL forms, the tree is congruent with the one in Fig. 1.

**Figure S2.** A maximum likelihood tree of the amino acid sequences of RbcL. All the sequences from the strain used in this study were included for constructing the tree with reference sequences which collected from published data (Badger and Bek, 2008; Prywes *et al*., 2023; Schulz *et al*., 2022). The tree was constructed using IQ-TREE with the LG+C60+G4+F substitution model and 1,000 ultrafast bootstrap replicates. Input sequences were trimmed using trimAl (-gt 0.8), resulting in 488 sequences with 466 amino acid positions. Bootstrap branch support values were calculated using BOOSTER (v0.1.2) from ultrafast bootstrap (1000 replicates). Unsupported branches with lower bootstrap values (≤90%) were collapsed, and branches with bootstrap values <95% were colored red. The scale bar indicates 1.0 changes per amino acid site.

## Notes

### Competing Interest Statement

The authors have declared no competing interest.

## References

1. Aono, R., Sato, T., Imanaka, T. and Atomi, H. 2015. A pentose bisphosphate pathway for nucleoside degradation in Archaea. Nat Chem Biol 11(5), 355–360.

2. Aono, R., Sato, T., Yano, A., Yoshida, S., Nishitani, Y., Miki, K., Imanaka, T. and Atomi, H. 2012. Enzymatic characterization of AMP phosphorylase and ribose-1,5-bisphosphate isomerase functioning in an archaeal AMP metabolic pathway. J Bacteriol 194(24), 6847–6855.

3. Badger, M.R. and Bek, E.J. 2008. Multiple Rubisco forms in proteobacteria: their functional significance in relation to CO2 acquisition by the CBB cycle. J Exp Bot 59(7), 1525–1541.

4. Banda, D.M., Pereira, J.H., Liu, A.K., Orr, D.J., Hammel, M., He, C., Parry, M.A.J., Carmo-Silva, E., Adams, P.D., Banfield, J.F. and Shih, P.M. 2020. Novel bacterial clade reveals origin of form I Rubisco. Nat Plants 6(9), 1158–1166.

5. Borrel, G., Adam, P.S. and Gribaldo, S. 2016. Methanogenesis and the Wood–Ljungdahl pathway: an Ancient, versatile, and fragile association. Genome Biology and Evolution 8(6), 1706–1711.

6. 6. Boutet, E., Lieberherr, D., Tognolli, M., Schneider, M. and Bairoch, A. (2007) UniProtKB/Swiss-Prot. Plant Bioinformatics: Methods and protocols. Edwards, D. (ed), pp. 89–112, Humana Press, Totowa, NJ.

7. Capella-Gutiérrez, S., Silla-Martínez, J.M. and Gabaldón, T. 2009. trimAl: a tool for automated alignment trimming in large-scale phylogenetic analyses. Bioinformatics 25(15), 1972–1973.

8. Da Costa, M.S., Rainey, F.A. and Nobre, M.F. (2006) The Prokaryotes: Volume 7: Proteobacteria: Delta, Epsilon Subclass. Dworkin, M., Falkow, S., Rosenberg, E., Schleifer, K.-H. and Stackebrandt, E. (eds), pp. 797–812, Springer New York, New York, NY.

9. Ferry, J.G., Smith, P.H. and Wolfe, R.S. 1974. *Methanospirillum*, a new genus of methanogenic bacteria, and characterization of *Methanospirillum hungatii* sp.nov. International Journal of Systematic and Evolutionary Microbiology 24(4), 465–469.

10. Field, C.B., Behrenfeld, M.J., Randerson, J.T. and Falkowski, P. 1998. Primary production of the biosphere: integrating terrestrial and oceanic components. Science 281(5374), 237–240.

11. Frolov, E.N., Kublanov, I.V., Toshchakov, S.V., Lunev, E.A., Pimenov, N.V., Bonch-Osmolovskaya, E.A., Lebedinsky, A.V. and Chernyh, N.A. 2019. Form III RubisCO-mediated transaldolase variant of the Calvin cycle in a chemolithoautotrophic bacterium. Proc Natl Acad Sci U S A 116(37), 18638–18646.

12. Fu, L., Niu, B., Zhu, Z., Wu, S. and Li, W. 2012. CD-HIT: accelerated for clustering the next-generation sequencing data. Bioinformatics 28(23), 3150–3152.

13. Garritano, A.N., Song, W. and Thomas, T. 2022. Carbon fixation pathways across the bacterial and archaeal tree of life. PNAS Nexus 1(5), pgac226.

14. 14. Imhoff, J.F. (2009) Phototrophic purple bacteria. In eLS, (Ed.).

15. Jones, R.M. and Johnson, D.B. 2015. *Acidithrix ferrooxidans* gen. nov., sp. nov.; a filamentous and obligately heterotrophic, acidophilic member of the *Actinobacteria* that catalyzes dissimilatory oxido-reduction of iron. Res Microbiol 166(2), 111–120.

16. Kalyaanamoorthy, S., Minh, B.Q., Wong, T.K.F., von Haeseler, A. and Jermiin, L.S. 2017. ModelFinder: fast model selection for accurate phylogenetic estimates. Nature Methods 14(6), 587–589.

17. Katoh, K., Kuma, K.-i., Toh, H. and Miyata, T. 2005. MAFFT version 5: improvement in accuracy of multiple sequence alignment. Nucleic Acids Research 33(2), 511–518.

18. Kishimoto, N., Fukaya, F., Inagaki, K., Sugio, T., Tanaka, H. and Tano, T. 1995. Distribution of bacteriochlorophyll a among aerobic and acidophilic bacteria and light-enhanced CO_2_-incorporation in *Acidiphilium rubrum*. FEMS Microbiology Ecology 16(4), 291–296.

19. Kono, T., Mehrotra, S., Endo, C., Kizu, N., Matusda, M., Kimura, H., Mizohata, E., Inoue, T., Hasunuma, T., Yokota, A., Matsumura, H. and Ashida, H. 2017. A RuBisCO-mediated carbon metabolic pathway in methanogenic archaea. Nat Commun 8, 14007.

20. Lartillot, N. and Philippe, H. 2006. Computing Bayes factors using thermodynamic integration. Systematic Biology 55(2), 195–207.

21. Liu, A.K., Kaeser, B., Chen, L., West-Roberts, J., Taylor-Kearney, L.J., Lavy, A., Gunzing, D., Li, W.J., Hammel, M., Nogales, E., Banfield, J.F. and Shih, P.M. 2023. Deep-branching evolutionary intermediates reveal structural origins of form I rubisco. Curr Biol 33(24), 5316–5325 e5313.

22. Mori, K., Kim, H., Kakegawa, T. and Hanada, S. 2003. A novel lineage of sulfate-reducing microorganisms: *Thermodesulfobiaceae* fam. nov., *Thermodesulfobium narugense*, gen. nov., sp. nov., a new thermophilic isolate from a hot spring. Extremophiles 7(4), 283–290.

23. Morris, R.L. and Schmidt, T.M. 2013. Shallow breathing: bacterial life at low O_2_. Nat Rev Microbiol 11(3), 205–212.

24. Müller, W.J., Tlalajoe, N., Cason, E.D., Litthauer, D., Reva, O., Brzuszkiewicz, E. and van Heerden, E. 2016. Whole genome comparison of *Thermus* sp. NMX2.A1 reveals principal carbon metabolism differences with closest relation *Thermus scotoductus* SA-01. G3 Genes|Genomes|Genetics 6(9), 2791–2797.

25. Nguyen, L.-T., Schmidt, H.A., von Haeseler, A. and Minh, B.Q. 2014. IQ-TREE: a fast and effective stochastic algorithm for estimating maximum-likelihood phylogenies. Molecular Biology and Evolution 32(1), 268–274.

26. Nishihara, A., Tsukatani, Y., Azai, C. and Nobu, M.K. 2024. Illuminating the coevolution of photosynthesis and Bacteria. Proc Natl Acad Sci U S A 121(25), e2322120121.

27. Norris, D.A.C.P.R. 1996. *Acidimicrobium ferrooxidans* gen. nov., sp. nov.: mixed-cultureferrous ironoxidation with *Sulfobacillus* species.

28. Prywes, N., Phillips, N.R., Tuck, O.T., Valentin-Alvarado, L.E. and Savage, D.F. 2023. Rubisco function, evolution, and engineering. Annu Rev Biochem 92, 385–410.

29. Sato, T., Atomi, H. and Imanaka, T. 2007. Archaeal type III RuBisCOs function in a pathway for AMP metabolism. Science 315(5814), 1003–1006.

30. Schulz, L., Guo, Z., Zarzycki, J., Steinchen, W., Schuller, J.M., Heimerl, T., Prinz, S., Mueller-Cajar, O., Erb, T.J. and Hochberg, G.K.A. 2022. Evolution of increased complexity and specificity at the dawn of form I Rubiscos. Science 378(6616), 155-+.

31. Seemann, T. 2014. Prokka: rapid prokaryotic genome annotation. Bioinformatics 30(14), 2068–2069.

32. Shiba, T. 1984. Utilization of light energy by the strictly aerobic bacterium *Erythrobacter* sp. OCH 114. The Journal of General and Applied Microbiology 30(3), 239–244.

33. Shiba, T. and Harashima, K. 1986. Aerobic photosynthetic bacteria. Microbiol Sci 3(12), 376–378.

34. Shimada, K. and Takaichi, S. (2024) Anoxygenic phototrophic bacteria. Elsevier, pp.111–138, 275–279

35. Shimizu, S., Ueno, A., Tamamura, S., Naganuma, T. and Kaneko, K. 2013. *Methanoculleus horonobensis* sp. nov., a methanogenic archaeon isolated from a deep diatomaceous shale formation. Int J Syst Evol Microbiol 63(Pt 11), 4320–4323.

36. Slobodkin, A.I., Slobodkina, G.B., Panteleeva, A.N., Chernyh, N.A., Novikov, A.A. and Bonch-Osmolovskaya, E.A. 2016. *Dissulfurimicrobium hydrothermale* gen. nov., sp. nov., a thermophilic, autotrophic, sulfur-disproportionating deltaproteobacterium isolated from a hydrothermal pond. Int J Syst Evol Microbiol 66(2), 1022–1026.

37. Sondergaard, D., Pedersen, C.N. and Greening, C. 2016. HydDB: A web tool for hydrogenase classification and analysis. Sci Rep 6, 34212.

38. Srinivas, T.N.R., Anil Kumar, P., Sasikala, C., Ramana, C.V. and Imhoff, J.F. 2007. *Rhodobacter vinaykumarii* sp. nov., a marine phototrophic alphaproteobacterium from tidal waters, and emended description of the genus *Rhodobacter*. Int J Syst Evol Microbiol 57(Pt 9), 1984–1987.

39. Tabita, F.R., Satagopan, S., Hanson, T.E., Kreel, N.E. and Scott, S.S. 2008. Distinct form I, II, III, and IV Rubisco proteins from the three kingdoms of life provide clues about Rubisco evolution and structure/function relationships. J Exp Bot 59(7), 1515–1524.

40. West-Roberts, J.A., Matheus-Carnevali, P.B., Schoelmerich, M.C., Al-Shayeb, B., Thomas, A.D., Sharrar, A., He, C., Chen, L.-X., Lavy, A., Keren, R., Amano, Y. and Banfield, J.F. 2021. The *Chloroflexi* supergroup is metabolically diverse and representatives have novel genes for non-photosynthesis based CO_2_ fixation.

41. Yurkov, V.V. and Beatty, J.T. 1998. Aerobic anoxygenic phototrophic bacteria. Microbiology and Molecular Biology Reviews 62(3), 695–724.

42. Yvenou, S., Allioux, M., Slobodkin, A., Slobodkina, G., Jebbar, M. and Alain, K. 2021. Genetic potential of *Dissulfurimicrobium hydrothermale*, an obligate sulfur-disproportionating thermophilic microorganism. Microorganisms 10(1).

43. Zhou, E.M., Xian, W.D., Mefferd, C.C., Thomas, S.C., Adegboruwa, A.L., Williams, N., Murugapiran, S.K., Dodsworth, J.A., Ganji, R., Li, M.M., Ding, Y.P., Liu, L., Woyke, T., Li, W.J. and Hedlund, B.P. 2018. *Thermus sediminis* sp. nov., a thiosulfate-oxidizing and arsenate-reducing organism isolated from Little Hot Creek in the Long Valley Caldera, California. Extremophiles 22(6), 983–991.

44. Zhou, Z., Tran, P.Q., Breister, A.M., Liu, Y., Kieft, K., Cowley, E.S., Karaoz, U. and Anantharaman, K. 2022. METABOLIC: high-throughput profiling of microbial genomes for functional traits, metabolism, biogeochemistry, and community-scale functional networks. Microbiome 10(1), 33.

