## Supplementary figures and images for "Exploring the diversity and physiological characteristics of RubisCO-mediated carbon fixation in culturable prokaryotes"

### Supplemental figures

Fig. S1 Nishihara et al.

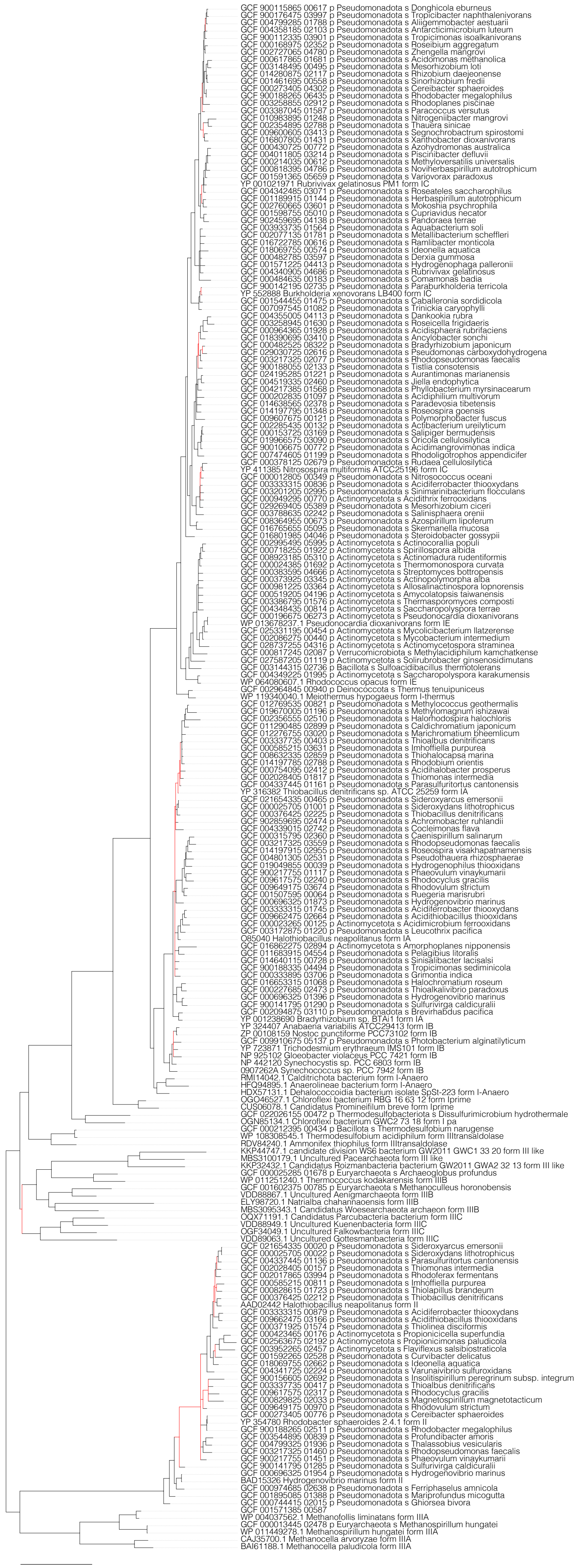

Fig. S2 Nishihara et al.

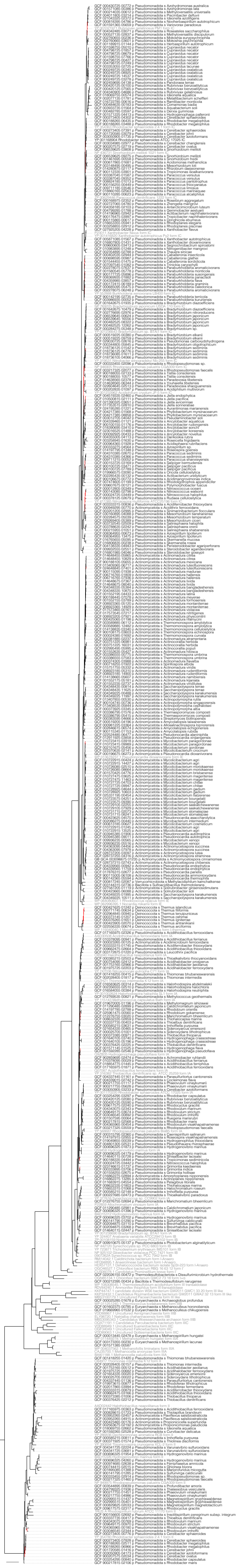
